# Live-cell Fluorescence Microscopy of HSV-1 Cellular Egress by Exocytosis

**DOI:** 10.1101/2023.02.27.530373

**Authors:** Melissa H. Bergeman, Michaella Q. Hernandez, Jenna Diefenderfer, Jake A. Drewes, Kimberly Velarde, Wesley M. Tierney, Junior A. Enow, Honor L. Glenn, Masmudur M. Rahman, Ian B. Hogue

## Abstract

1.1

The human pathogen Herpes Simplex Virus 1 (HSV-1) produces a lifelong infection in the majority of the world’s population. While the generalities of alpha herpesvirus assembly and egress pathways are known, the precise molecular and spatiotemporal details remain unclear. In order to study this aspect of HSV-1 infection, we engineered a recombinant HSV-1 strain expressing a pH-sensitive reporter, gM-pHluorin. Using a variety of fluorescent microscopy modalities, we can detect individual virus particles undergoing intracellular transport and exocytosis at the plasma membrane. We show that particles exit from epithelial cells individually, not bulk release of many particles at once, as has been reported for other viruses. In multiple cell types, HSV-1 particles accumulate over time at the cell periphery and cell-cell contacts. We show that this accumulation effect is the result of individual particles undergoing exocytosis at preferential sites and that these egress sites can contribute to cell-cell spread. We also show that the viral membrane proteins gE, gI, and US9, which have important functions in intracellular transport in neurons, are not required for preferential egress and clustering in non-neuronal cells. Importantly, by comparing HSV-1 to a related alpha herpesvirus, pseudorabies virus, we show that this preferential exocytosis and clustering effect is cell type-dependent, not virus dependent. This preferential egress and clustering appears to be the result of the arrangement of the microtubule cytoskeleton, as virus particles co-accumulate at the same cell protrusions as an exogenous plus end-directed kinesin motor.

**Importance:** Alpha herpesviruses produce lifelong infections of their human and animal hosts. The majority of people in the world are infected with Herpes Simplex Virus 1 (HSV-1), which typically causes recurrent oral or genital lesions. However, HSV-1 can also spread to the central nervous system, causing severe encephalitis, and might also contribute to the development of neurodegenerative diseases. Many of the steps of how these viruses infect and replicate inside host cells are known in depth, but the final step, exiting from the infected cell, is not fully understood. In this study, we engineered a novel variant of HSV-1 that allows us to visualize how individual virus particles exit from infected cells. With this imaging assay, we investigated preferential egress site formation in certain cell types and their contribution to cell-cell spread of HSV-1.

## 1.2 Introduction

The human pathogen, Herpes Simplex Virus type 1 (HSV-1), is a member of the alpha herpesvirus sub-family, which includes several endemic human pathogens, economically-important veterinary pathogens, and zoonotic pathogens that can be severely neuroinvasive. The generalities of alpha herpesvirus assembly and egress pathways are known, but the spatiotemporal details remain unclear. Viral DNA replication and packaging occurs in the nucleus, nuclear egress occurs by transient envelopment/de-envelopment at the nuclear membranes [1–3], and secondary envelopment occurs on intracellular membranes derived from the secretory and endocytic pathways [4–11]. Following secondary envelopment, the secretory organelle containing the enveloped virion traffics to the plasma membrane, predominantly using microtubule motors, where it is released by exocytosis [12–17]. However, the precise molecular details, virus-host interactions, and dynamics of this process are not fully understood. While fluorescence microscopy has been widely used to determine the relationships between viral proteins and cellular markers, it lacks the spatial resolution to determine the precise assembly state of the virion. Electron microscopy has also provided many insightful results, but this technique is confounded by difficulties in sample preparation, small sample sizes, and the fact that samples are fixed and static [18–21]. This last constraint is of particular concern as it does not offer insights into the dynamic aspects of the viral replication cycle in infected cells. To overcome some of these limitations, we previously developed a live-cell fluorescence microscopy method to study exocytosis of the important veterinary and zoonotic virus, Pseudorabies Virus (PRV; suid alphaherpesvirus 1) [16, 17]. In these studies, we showed that PRV particles exit from infected cells by exocytosis using cellular secretory mechanisms, are mainly released as single particles from individual secretory vesicles, and the spatial distribution viral exocytosis is largely uniform across the adherent cell surface (in PK15 cells, a porcine kidney epithelial cell line). However, other studies focusing on HSV-1 showed that viral proteins and particles form large clusters at particular locations on the plasma membrane, which the authors inferred was the result of viral exocytosis at preferential sites (in Vero cells, an African green monkey kidney epithelial cell line) [22–25].

A variety of egress modes have been observed with other viruses: The beta herpesvirus, human cytomegalovirus (HCMV), was recently shown to exit by bulk release – exocytosis of many particles from a larger organelle – in human foreskin fibroblast (HFF-1) cells [26]. Both flaviviruses and coronaviruses have been observed by electron microscopy to accumulate large numbers of virus particles in large intracellular organelles, but it is unclear whether these large organelles mediate bulk release or if there are subsequent intracellular sorting steps to release single virions from individual exocytosis events [27, 28]. In retroviruses, HIV-1 has been observed to assemble and exit preferentially at the trailing uropod of polarized T cells [29, 30], and human T-lymphotropic virus (HTLV) forms large accumulations of virions and extracellular matrix (termed “viral biofilms”) on the cell surface, which may promote more efficient cell-cell spread [31–34]. Thus, the relationship between exocytosis (single particles in individual secretory vesicles, versus bulk release of many particles from a larger organelle) and accumulation at preferential locations on the cell surface following exocytosis varies according to the particular virus and cell type. But, these features of viral egress are likely important for subsequent cell-cell spread.

In the present study, we have extended our previous work on PRV to study the egress of HSV-1 particles via live-cell fluorescence microscopy. To construct a model system allowing us to visualize HSV-1 exocytosis, we engineered a recombinant strain of HSV-1 that expresses superecliptic pHluorin on an extravirion loop of the multipass transmembrane glycoprotein M (gM-pHluorin). A variant of GFP, pHluorin was developed as a means to image secretory vesicle exocytosis in a variety of cell types, including neurons [35, 36]. Following secondary envelopment, pHluorin is quenched in the acidic lumen of secretory vesicles (pH of 5.2-5.7) [36, 37]. When the secretory vesicle fuses with the plasma membrane to release the virus particle to the extracellular medium (pH ⁓7.5), pHluorin is dequenched and becomes brightly fluorescent, allowing the unambiguous identification of individual viral exocytosis events [16, 17, 37, 38] (Figure 1A).

**Figure 1.**
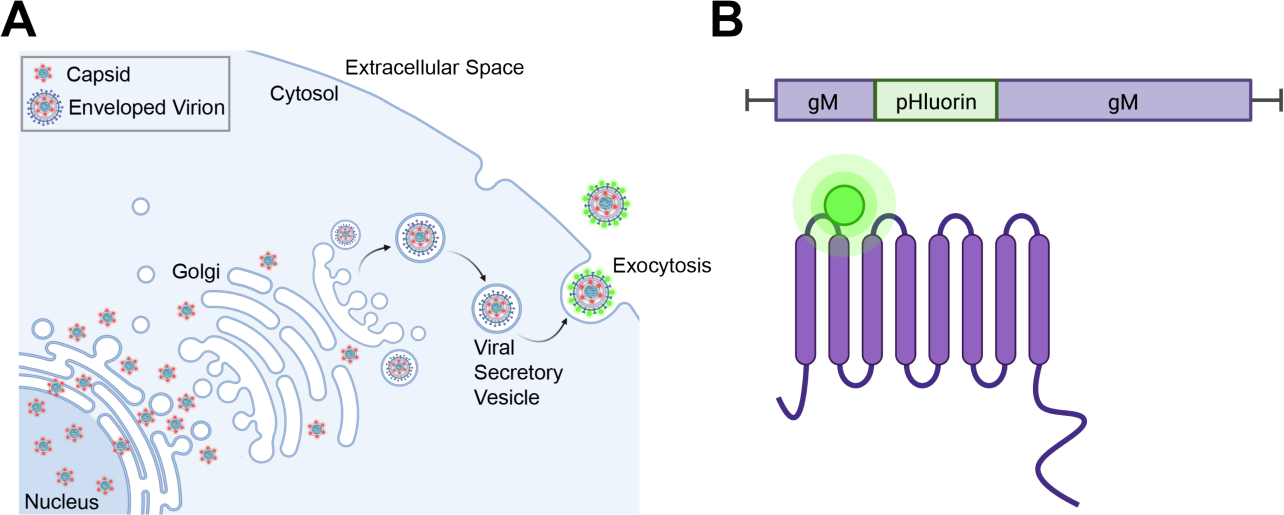
HSV-1 Egress and gM-pHluorin Insert. **A.** Schematic of HSV-1 egress from infected cells. Nonenveloped virus capsids (red) exit the nucleus and traffic to the site of secondary envelopment. Following secondary envelopment, virions are transported to the plasma membrane by acidic secretory organelles. pHluorin (green) dequenches upon exocytosis at the plasma membrane. **B.** Schematic of gM-pHluorin, with the pHluorin moiety (green) inserted into the first extravirion loop of gM (purple).

Using this technique, we show that HSV-1 exits from infected cells by exocytosis of individual virus particles, not bulk release of many virions at once. In some cell types, viral exocytosis occurs at preferential plasma membrane sites, leading to the gradual accumulation of large clusters of virus particles, but we show that this phenomenon is cell-type dependent. Consistent with previous reports [22], mutations in viral membrane proteins gE, gI, and US9 were not essential for preferential viral egress and accumulation into clusters. To characterize the cellular mechanisms responsible for this phenomenon, we show that an exogenous kinesin microtubule motor co-accumulates at sites of cluster formation, indicating that the arrangement of the microtubule cytoskeleton likely directs virus particle transport to particular locations, resulting in preferential egress and cluster formation at these sites. Finally, using timelapse confocal imaging, we show that these large peripheral accumulations of virus particles form at sites of cell-cell contact and contribute to cell-cell spread of infection.

## 1.3 Results

### Insertion of pHluorin into gM

To produce the recombinant strain HSV-1 IH01, we inserted the pHluorin coding sequence into the gM (UL10) gene in the HSV-1 genome by homologous recombination between a synthesized shuttle plasmid and purified HSV-1 DNA. The construct was designed to insert the pHluorin moiety into the first extravirion/lumenal loop of gM (Figure 1B). A second recombinant, HSV-1 IH02, expressing gM-pHluorin and an mRFP-VP26 capsid tag, was produced by co-infecting HSV-1 IH01 and HSV-1 OK14 [39], and purifying two-color plaques. We confirmed the correct recombination occurred by PCR amplification and Sanger sequencing (Supplemental Material 1), and also the expression of gM-pHluorin by western blot of infected cell lysates. The western blots were probed with anti-gM and anti-GFP antibodies simultaneously (Figure 2A). Viral membrane proteins frequently produce complex banding patterns due to post-translational modifications like glycosylation and aggregation of these highly hydrophobic proteins during sample prep [40, 41]. Cells infected with parental strains HSV-1 17syn^+^ and OK14 produced major gM-immunoreactive bands near the predicted 51 kDa of native gM, whereas cells infected with the recombinant HSV-1 IH01 and IH02 strains produced bands that are immunoreactive to both gM and GFP antibodies, and shifted ⁓30 kDa, consistent with the predicted gM-pHluorin fusion (Figure 2A).

**Figure 2.**
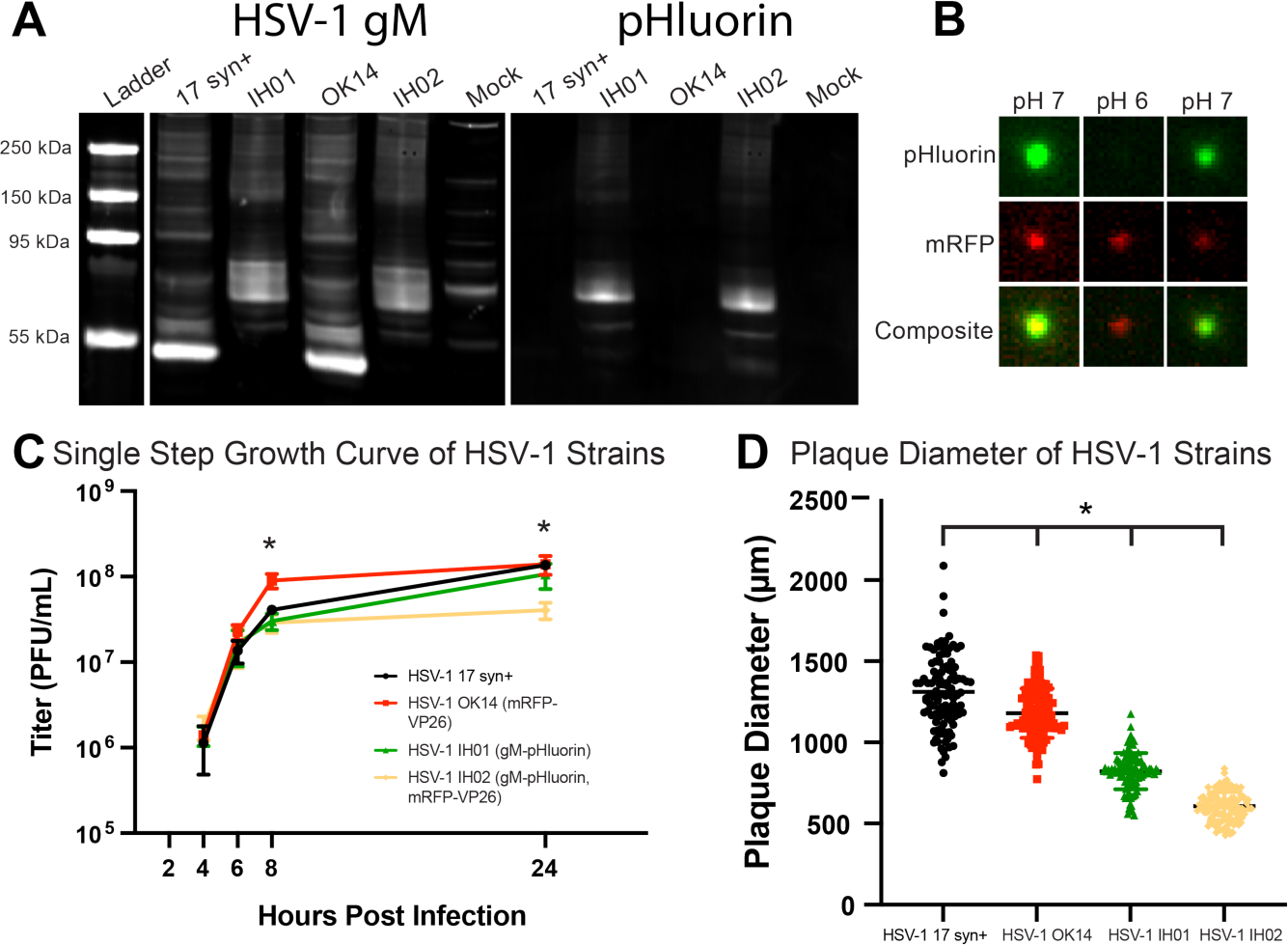
pHluorin Expression and Impact on Viral Replication **A.** Western blot detecting HSV-1 gM and pHluorin. Vero cells were infected with HSV-1 17syn^+^, IH01, OK14, and IH02, or mock-infected. Blots were probed with primary antibodies detecting gM and pHluorin, and imaged using fluorescent secondary antibodies to detect HSV-1 gM and pHluorin simultaneously. **B.** gM-pHluorin is incorporated into virus particles, and exhibits reversibly pH-sensitive fluorescence. Freshly-prepared supernatants from Vero cells infected with HSV-1 IH02 were spotted onto glass bottom dishes. Particles were imaged at pH ⁓7, pH ⁓6, and pH ⁓7. A representative virus particle is shown with gM-pHluorin and mRFP-VP26. Images represent 3.6×3.6 *µ*m. **C.** Single Step Growth Curve. Vero cells were infected with HSV-1 17syn^+^, OK14, IH01, or IH02 and harvested at 4, 6, 8, and 24 hpi. Infections were done in triplicate for each virus at each time point. Samples were titered by serial-dilution plaque assay. Error bars represent standard deviation, asterisk represents statistical significance with p<0.05. Means of each time point were compared with one-way ANOVA in Graphpad Prism. **D.** Plaque size measurements. At 4 days post-infection, virus plaques were imaged, and the zone of clearance diameter was measured in Fiji software and used to calculate mean plaque sizes (n=100). Asterisk indicates statistical significance with p<0.05. HSV-17 was compared with each fluorescent strain with a Student’s T-test.

### gM-pHluorin Labels Virus Particles and Exhibits pH-Sensitive Fluorescence

To determine whether gM-pHluorin is incorporated into individual virus particles, we spotted ⁓100*µ*l of freshly-prepared infected cell supernatants onto a glass coverslip, and imaged by fluorescence microscopy (Figure 2B). To measure the pH sensitivity of the gM-pHluorin fluorescence, we added an excess of PBS buffer at pH ⁓6 followed by an excess of PBS buffer at pH ⁓7. gM-pHluorin incorporated into virus particles exhibited reversible pH-dependent green fluorescence, whereas the mRFP-VP26 capsid tag exhibited a non-pH-sensitive reduction in fluorescence due to photobleaching (Figure 2B).

### Virus Replication

To determine whether the recombinant HSV-1 IH01 and IH02 strains replicate comparably to the parental viruses, we performed single-step growth curves (Figure 2C) and measured plaque size (Figure 2D) on Vero cell monolayers. Compared to the parental HSV-1 17syn^+^ strain, HSV-1 OK14, IH01, and IH02 exhibited a modest delay in replication at 8 hours post-infection (hpi). However, by 24 hpi, HSV-1 OK14 and IH01 had titers equivalent to 17syn^+^, but HSV-1 IH02 exhibited a modest <1 log defect (Figure 2C). These data suggest that the gM-pHluorin and mRFP-VP26 fluorescent protein fusions result in a small reduction in viral replication. Consistent with these results, plaque sizes of the HSV-1 OK14 and IH01 were also reduced (Figure 2D). Although these defects could also be explained by other mutations arising during construction, we initially characterized two different recombinants and several independent clones of each, with similar results.

### Live-Cell Fluorescence Microscopy of Virus Particle Exocytosis

To investigate virus particle exocytosis with our model system, we infected Vero cells with HSV-1 IH01 at a high multiplicity of infection (MOI) to roughly synchronize viral infection, and imaged by live-cell fluorescence microscopy at approximately 5-6 hpi. This time point represents the earliest production of viral progeny, prior to the onset of cytopathic effects (CPE). To compare to our previous studies of PRV [16], we also infected PK15 cells with PRV 483, which expresses orthologous gM-pHluorin and mRFP-VP26 fusions. We identified productively infected cells by imaging in widefield fluorescence mode to detect mRFP-VP26 fluorescence in the nucleus and gM-pHluorin green fluorescence on the plasma membrane and in intracellular membranes. We then acquired timelapse movies in Total Internal Reflection Fluorescence (TIRF) microscopy mode, which excludes out-of-focus fluorescence and emphasizes particle dynamics near the adherent cell surface (Figure 3A).

**Figure 3.**
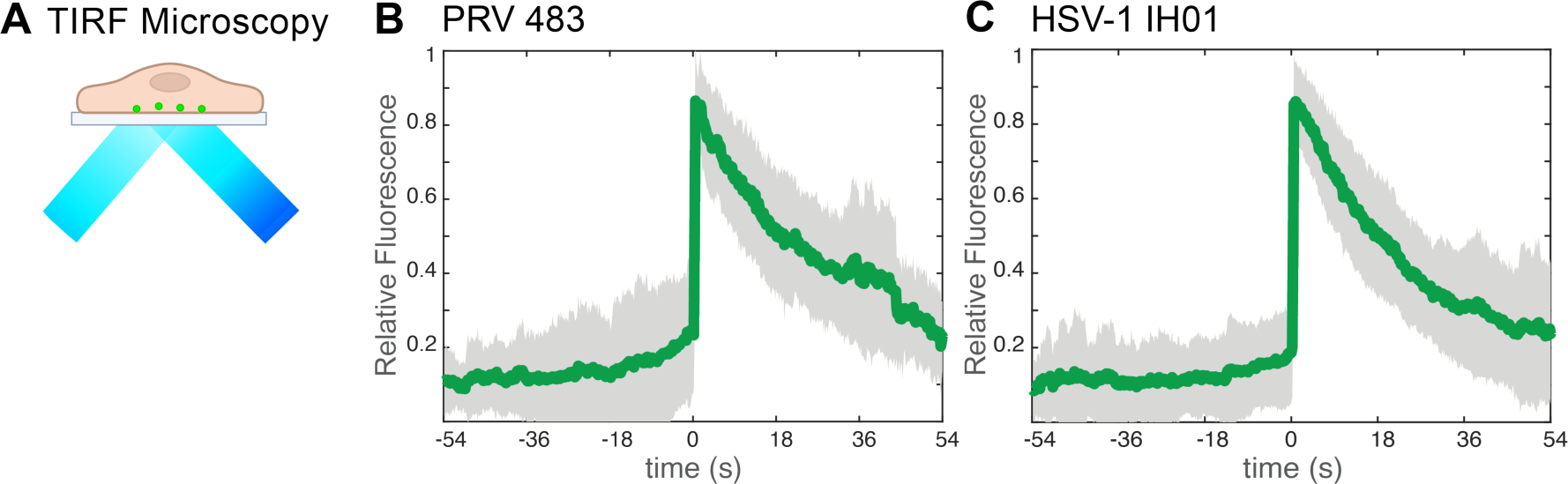
TIRF Microscopy of HSV-1 Exocytosis **A**. Schematic of TIRF microscopy. The excitation laser excites fluorescent molecules near the coverslip, and excludes out-of-focus fluorescence from deeper in the cell. **B-C**. Relative fluorescence intensity of gM-pHluorin before, during, and after exocytosis of individual virus particles. Green line represents mean fluorescence and gray shading indicates standard deviation. **B** PRV 483 exocytosis events in PK15 cells, at 4-5 hpi (n=31). **C** HSV-1 IH01 exocytosis events in Vero cells, at 5-6 hpi (n=67).

As previously reported with PRV, virus particle exocytosis events are characterized by the sudden (<90 ms) appearance of green gM-pHluorin fluorescence, which then remains pucntate and mostly immobile during the time of imaging (>2-3 min) [16, 17]. Exocytosis events that do not contain a particle, characterized by rapid diffusion of gM-pHluorin into the plasma membrane, are excluded from this analysis.. To quantify this process over many exocytosis events, we measured the relative fluorescence intensity at exocytosis sites for 54 sec before and after each exocytosis event, aligned all data series to a common time=0, and calculated the ensemble average over many events (Figure 3B-C). Prior to exocytosis at time=0, the relative gM-pHluorin fluorescence remains low, consistent with pHluorin quenching in the acidic lumen of the viral secretory vesicle. At the moment of exocytosis, gM-pHluorin fluorescence increases suddenly due to dequenching at extracellular pH. Finally, the fluorescence decays gradually, which represents a combination of: 1. diffusion of gM-pHluorin that is incorporated into the vesicle membrane; 2. occasional movement of the cell or virus particle after exocytosis; 3. photobleaching. In total, we quantified 67 HSV-1 IH01 exocytosis events from over 30 individual Vero cells across six replicate experiments. These data are consistent with our previous studies of PRV exocytosis [16, 17], validating that this approach works for HSV-1.

### HSV-1 Particles Accumulate at Preferential Exocytosis Sites in Multiple Cell Types

Previous studies showed that HSV-1 structural proteins and particles accumulate in large clusters at the adherent edges of Vero cells and cell-cell junctions in epithelial cells [22–25]. However, based on static fluorescence and electron microscopy images, it is unclear if virus particles gradually accumulate in these clusters due to preferential exocytosis at these sites, if large clusters are deposited at once due to bulk release (as recently observed with HCMV [26]), or if virus particles accumulate in clusters later in infection due to cell movement and rounding associated with cytopathic effects. Previously, we did not observe preferential exocytosis sites or large clusters of virus particles with PRV in PK15 cells [16, 17], so it was unclear whether this represents a difference in virus biology or that of host cell biology.

To better understand how these large clusters of virus particles form, we infected Vero or PK15 cells with HSV-1 IH02 and imaged at 6-7 hpi. At this time point, HSV-1 IH02 particles were beginning to accumulate in peripheral clusters in Vero cells (Figure 4A). A representative time course is provided in supplemental material, and illustrated using a maximum difference projection (Figure 4B, Movie S1). Maximum difference projections show where fluorescence intensity increases most rapidly, which emphasizes exocytosis events and particle movement, and deemphasizes static features that do not change during the course of imaging. In this representative Vero cell with multiple exocytosis events over time (4:01 min:sec), we observed exocytosis of particles containing green gM-pHluorin only, which we infer to be non-infectious L-particles (green boxes), and particles containing both gM-pHluorin and mRFP-VP26 capsids, which we infer to be virions (yellow circles). The diameter of detected fluorescence in these exocytosis events is consistent with diffraction-limited HSV-1 particles. Some viral exocytosis events appeared to be clustered near the cell periphery and cell protrusions, suggesting that the large peripheral clusters that appear later in infection accumulate gradually by the exocytosis of individual particles (Figure 4B, Movie S1). In this representative exocytosis event, a virus particle (pseudocolored magenta) arrives at the location of exocytosis, is largely immobile for 10 s, and then green pHluorin dequenches, indicating exocytosis (Figure 4B, right panel)

**Figure 4.**
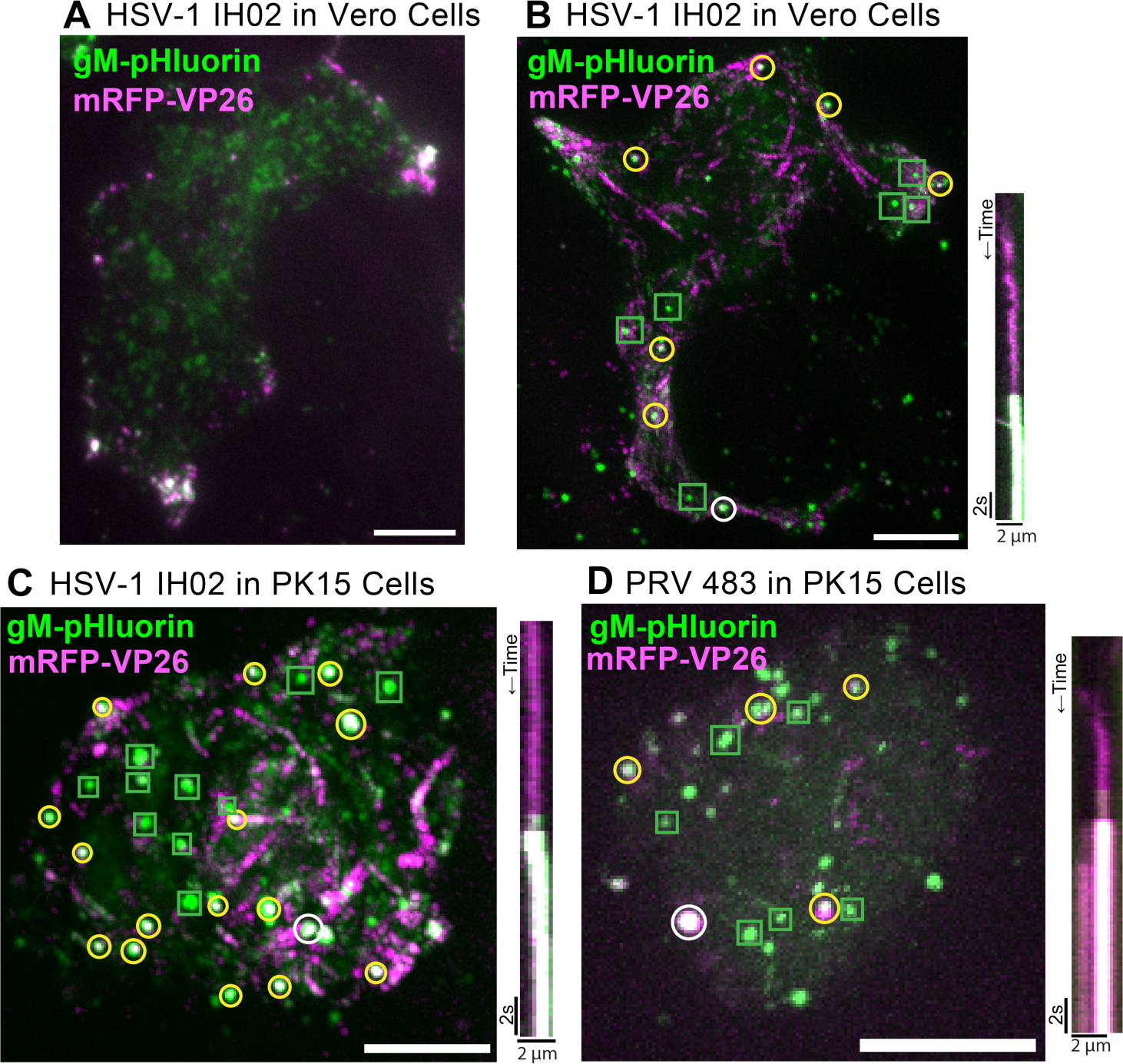
Single Virus Particle Exocytosis of HSV-1 and PRV **A**. HSV-1 IH02 infection in Vero cells, imaged by TIRF microscopy. Virus capsids (magenta) and gM-pHluorin (green) accumulate in clusters at the cell periphery at 6-7 hpi. **B-D** Maximum difference projections showing viral exocytosis events over time (left image). Green squares indicate exocytosis events containing gM-pHluorin (L-particles), yellow circles indicate exocytosis events containing both gM-pHluorin and mRFP-VP26 (virions), and white circles indicate the exocytosis events shown in the accompanying kymographs (right image). **B**. HSV-1 IH02 in Vero cells. Projection image represents 4:01 min:sec of imaging time at 6-7 hpi. Movie S1 shows these exocytosis events over this time course. **C**. HSV-1 IH02 in PK15 cells. Projection image represents 3:56 min:sec of imaging time at 6-7 hpi. Movie S2 shows these exocytosis events over this time course. **D**. PRV 483 in PK15 cells. Projection image represents 1:45 min:sec of imaging time at 4-5 hpi. In all panels, scale bars represent 10*µ*m. Images are representative of hundreds of cells over >10 independent experimental replicates.

Because we previously reported no such accumulations of virus particles with PRV in PK15 cells [16, 17], we compared HSV-1 IH02 to PRV 483 in PK15 cells (Figure 4C-D, Movie S2). In contrast to Vero cells, there appear to be no large accumulations and no preferential sites of HSV-1 egress in PK15 cells (Figure 4C, Movie S2), similarly to PRV in PK15 cells (Figure 4D). Exocytosis events in the representative PK15 cell (Figure 4C, Movie S2) are distributed across most of the footprint of the cell, and both individual virions and L-particles are easily distinguishable even with the many exocytosis events occurring during this time course (3:56 min:sec).

To determine whether this clustering occurs in a more biologically-relevant primary cell type, we prepared rat embryonic fibroblasts (REFs), and infected them with HSV-1 IH02 or PRV 483. In these cells, it was difficult to assess whether virus particles clustered to the same degree as in Vero cells, because REFs exhibited cytopathic effects (CPE) earlier than in the transformed cell lines; however, peripheral accumulations were observed prior to significant rounding and CPE (Figure 5A). Similarly to Vero and PK15 cells, HSV-1 particles appeared to exit from REFs in the form of individual exocytosis events (Figure 5B-C), again suggesting that the observed peripheral clusters accumulate gradually over time.

**Figure 5.**
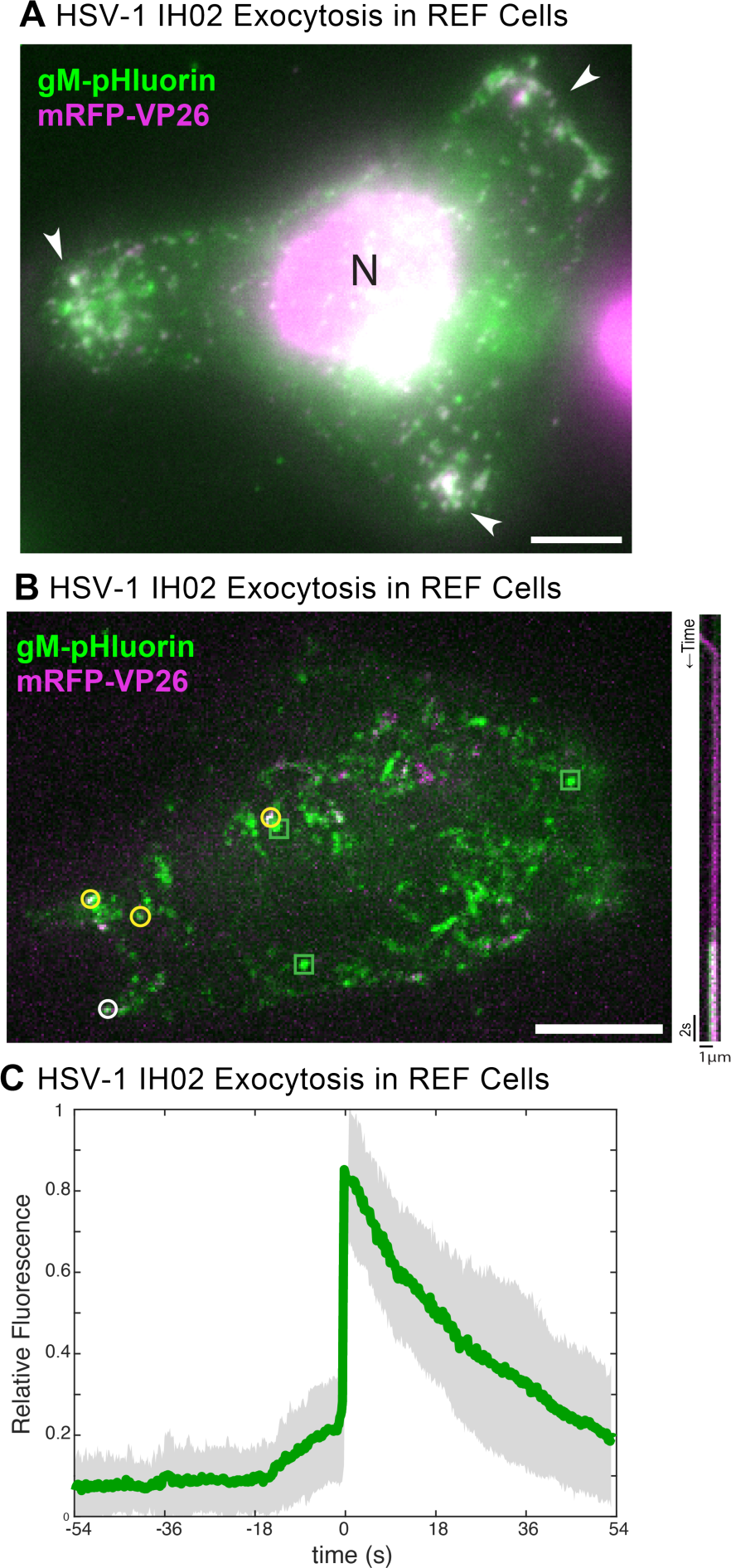
HSV-1 Exocytosis in Primary REF Cells **A**. HSV-1 IH02 infection in REF cells, with peripheral clusters of virus particles (arrows). Image acquired via widefield fluorescence to detect the cell nucleus (N). **B**. Maximum difference projections showing viral exocytosis events over time (left image), of HSV-1 IH02 in REF cells. Green squares indicate exocytosis events containing gM-pHluorin (L-particles), yellow circles indicate exocytosis events containing both gM-pHluorin and mRFP-VP26 (virions), and white circle indicates the exocytosis event shown in the accompanying kymograph (right image). **A-B**. Images are representative of approximately 10 cells from 3 experimental replicates. **C**. Relative fluorescence intensity of gM-pHluorin before, during, and after exocytosis of individual virus particles of HSV-1 IH02 in REF cells. Green line represents mean fluorescence and gray shading indicates standard deviation (n=42).

Because HSV-1 co-evolved in humans, we also investigated the clustering egress phenotype in human-derived cell lines: Panc-1 cells, a human pancreatic epithelioid sarcoma cell line, and a derivative of Panc-1 cells with stable RNAi knockdown of the cellular RNA helicase DHX9. The cellular functions of DHX9 remain enigmatic, but it is involved in epithelial-mesenchymal transition (EMT) in cancer biology, and has been shown to function as an antiviral factor [42]. In antiviral signaling, DHX9 expression is upregulated by IL-1 and TNF signaling, and, in turn, promotes NF-*κ*B and JAK-STAT signaling [43–45].

We infected Panc-1 and DHX9 knockdown cells with a high MOI of HSV-1 IH02, and imaged at 6-7 hpi. Consistent with the role of DHX9 in EMT, the parental Panc-1 cells exhibited a rounded cobblestone epithelial morphology (Figure 6A) [46–49], whereas the DHX9 knockdown cells exhibited elongated cell extensions typical of mesenchymal cell morphology (Figure 6B) [46]. Also, consistent with the role of DHX9 as an antiviral signaling factor, DHX9 knockdown cells appeared to be more permissive to HSV-1 infection, with greater expression of gM-pHluorin and mRFP-VP26 structural proteins in our microscopy assays (Figure 6A-B). We readily observed individual viral exocytosis events (Figure 6B-C), and peripheral accumulations of virus particles (Figure 6B,D), similarly to Vero cells. There appeared to be slightly greater accumulation/clustering of virus particles in DHX9 knockdown cells (Figure 6A,B,D), which may be due to alterations in cytoskeleton and cell morphology related to DHX9 function in EMT, or greater viral replication and expression of viral structural proteins related to DHX9 antiviral functions. While the percentage of cells exhibiting clustering was not significantly different (p>0.05; Figure 6D), in cells that did exhibit clustering, there appeared to be slightly greater accumulation of virus particles in DHX9 knockdown cells (Figure 6A,B). This may be due to alterations in cytoskeleton and cell morphology related to DHX9 function in EMT, or greater viral replication and expression of viral structural proteins related to DHX9 antiviral functions.

**Figure 6.**
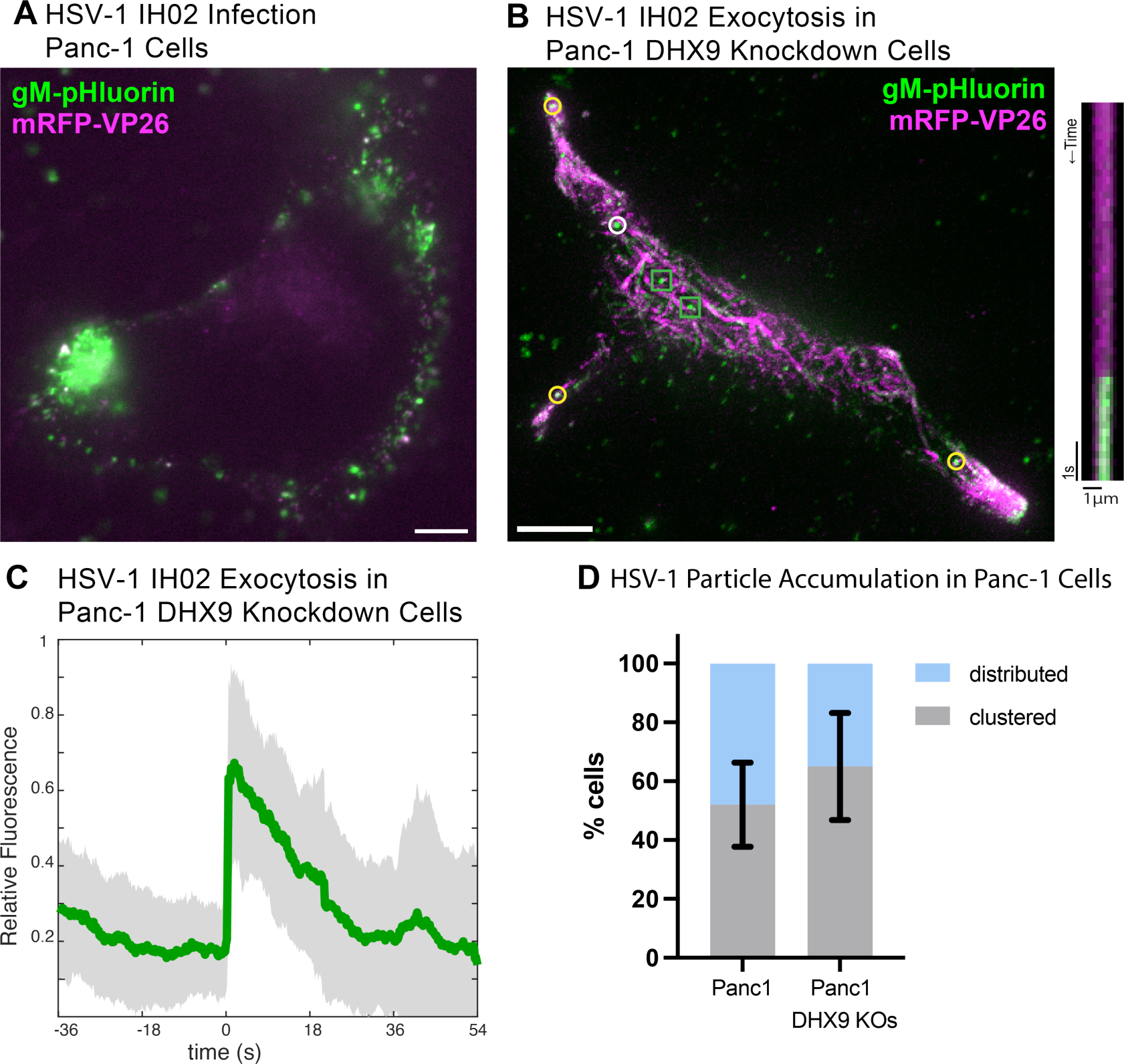
HSV-1 Exocytosis in Human Panc-1 Cells **A-B**. HSV-1 IH01 infection in human Panc-1 cells versus DHX9 knockdown cells. Compared to parental Panc-1 cells, DHX9 knockdown cells exhibited more of an elongated mesenchymal morphology, greater expression of viral structural proteins, and greater clustering at cell extensions, as assessed by fluorescence microscopy. Images are representative of 10 cells in 3 experimental replicates. Scale bars represent 10*µ*m. **C**. Relative fluorescence intensity of gM-pHluorin before, during, and after exocytosis of individual virus particles of HSV-1 IH02, in DHX9 knockdown cells. Green line represents mean fluorescence and gray shading indicates standard deviation (n=13). **D**. Percentage of imaged cells with a clustered or distributed virus particle distribution during infection (n=100). Not significantly different (p>0.05 by Student’s T test). Error bars represent standard deviation.

### Viral Membrane Proteins gE, gI, and US9 are Not Required for Clustered Egress in Vero and REF Cells

The three viral membrane proteins, gE, gI, and US9, have important functions in both HSV-1 and PRV egress. The clinical isolate HSV-1 MacIntyre and attenuated vaccine strain PRV Bartha, spread only in the retrograde direction and are incapable of anterograde spread in host nervous systems due mutations that disrupt the gE, gI, and US9 genes [50–55]. These proteins contribute to secondary envelopment, recruit microtubule motors for particle transport in multiple cell types [14, 56–58], and are required for axonal sorting and anterograde axonal spread in neurons [23, 50, 59–62]. However, it is not clear how mutations in gE, gI and US9 might affect egress in non-neuronal cells.

HSV-1 OK14, which is based on the 17syn^+^ laboratory strain, expresses functional gE, gI, and US9 [39, 63]. HSV-1 425 is based on the HSV-1 MacIntyre strain, which contains many polymorphisms, including mutations that disrupt gE/gI/US9 function [54, 55]. Both viruses express an mRFP-VP26 capsid tag. At about 5 hpi, we manually categorized infected cells in random fields of view based on the presence of virus particle clusters at the cell periphery. Cells infected with HSV-1 425 demonstrated a roughly similar proportion of clustering compared to HSV-1 OK14 in Vero cells (Figure 7B). These results show that gE, gI, and US9 are not strictly required for clustered egress of HSV-1. Because HSV-1 MacIntyre contains many polymorphisms compared to 17syn^+^, there remains a possibility that MacIntyre contains compensatory mutations that allow for a clustered egress phenotype in the absence of gE/gI/US9 function; however, these results are consistent with previous work by Mingo, et al. [22], who showed that gE is not necessary for cluster formation in Vero cells.

**Figure 7.**
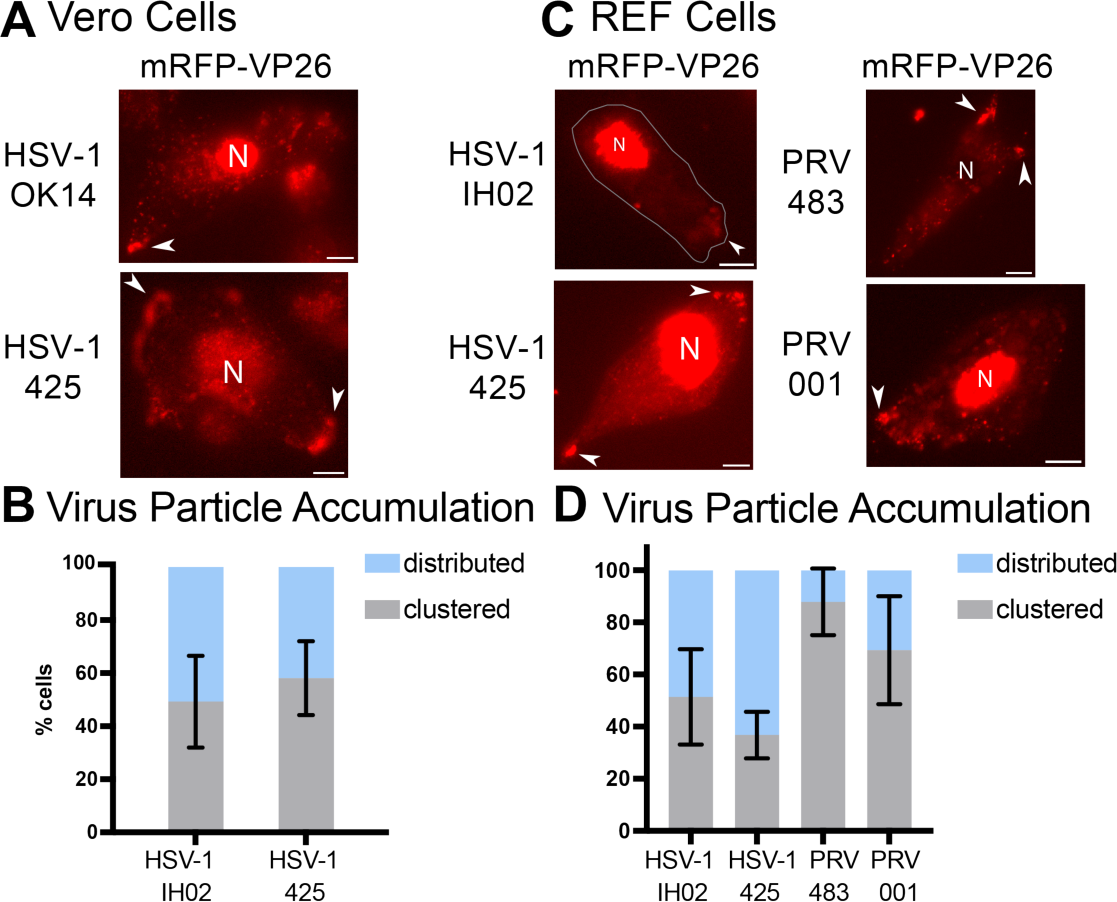
Viral Membrane Proteins gE, gI, & US9 Are Not Required for Peripheral Clustering **A-B**. HSV-1 OK14 (gE/gI/US9 wild-type) and HSV-1 425 (gE/gI/US9 null) viruses form peripheral accumulations of particles in Vero cells at 6-7 hpi. C-D. HSV-1 IH02 (gE/gI/US9 wild-type), HSV-1 425, PRV 483 (gE/gI/US9 wild-type), and PRV IH001 (gE/gI/US9 null) viruses form accumulations in primary REF cells 6-7 hpi. Nuclei (N) and virus particle clusters (arrows) are indicated. All scale bars represent 10*µ*m. **B-D**. Percentage of imaged cells that showed a distributed or clustered phenotype during infection (n=100) across 3 independent experiments. Error bars represent standard deviation.

To compare PRV to HSV-1, and further assess the function of gE, gI, and US9 in isogenic virus strains, we also infected cells with PRV recombinants. PRV 483 is based on the common Becker laboratory strain, which expresses functional gE, gI, and US9 [64, 65]. We also constructed PRV IH001, which contains the PRV Bartha vaccine strain deletion that removes the gE/gI/US9 genes, in an isogenic Becker genetic background. Both of these viruses express gM-pHluorin and mRFP-VP26. While PRV will infect and form plaques on Vero cells, we were unable to achieve sufficient levels of infection for our microscopy experiments - it is possible that the efficiency of plating of PRV in Vero cells is too low to achieve a high-MOI roughly synchronous infection in our experimental conditions. To overcome this limitation, we instead infected primary REF cells, which support robust infection of both viruses. HSV-1 and PRV exhibited clustering at the cell periphery, and gE, gI, and US9 proteins were not required for clustering. These results further reinforce the idea that this clustering effect is common to both HSV-1 and PRV, and varies by cell type. However, the polarized trafficking that is mediated by gE/gI/US9 in neurons is not essential for clustered egress in these non-neuronal cell types.

### Exogenous Kinesin Microtubule Motors Co-accumulate with Virus Particles in Peripheral Clusters

To better characterize the peripheral accumulations of virus particles we observed, we identified a cellular marker that labels the “corners” and tips of cell extensions before and during virus replication and egress. A large body of literature has shown that microtubule motors mediate intracellular transport of HSV-1 particles [12, 66–69]. Based on our observations above, we hypothesized that the arrangement of the microtubule cytoskeleton might explain why virus particles accumulate at particular subcellular locations during egress. We reasoned that microtubules may be preferentially arranged with their (+) ends at these peripheral sites, leading to the preferential transport to and egress at these sites.

In preliminary attempts, we were unable to adequately resolve individual microtubules in live cell microscopy due to the high abundance of microtubules and tubulin protein throughout the cell. While microtubule (+) end binding proteins (e.g. EB1, EB3) can be used to mark the growing (+) ends of dynamic microtubules, they do not effectively mark the (+) ends of stabilized microtubules, and HSV-1 infection promotes microtubule stabilization [70–72]. Therefore, as a probe of microtubule arrangement, we expressed KIF1A-EmGFP, an exogenous (+) end-directed microtubule motor that accumulates at microtubule (+) ends in live cells. KIF1A is a kinesin-3 family motor that is highly expressed in neurons, where it contributes to axonal sorting and transport of cellular cargoes and virus particles. However, KIF1A is not strongly expressed in most non-neuronal cells, including Vero cells [73]. We infected Vero cells with HSV-1 OK14 and an amplicon vector expressing KIF1A-EmGFP. At 6-7 hpi, we identified co-infected cells by widefield fluorescence microscopy, and found that KIF1A-EmGFP and virus particles co-accumulated at the tips of cell extensions (Figure 8A, arrows). Figure 8A shows three representative cells: In all three cells, mRFP-VP26 is visible in the nucleus, indicating active viral replication and capsid assembly. In the top cell, KIF1A-EmGFP and virus particles are strongly accumulated at the left and right ends of this elongated cell. In the middle cell, there are some peripheral accumulations of virus particles, but this cell is not expressing appreciable amounts of KIF1A-EmGFP. In the bottom cell, KIF1A-EmGFP is accumulated at the tips of cell extensions, but mRFP-VP26 is largely restricted to the nucleus, indicating that peripheral accumulations of KIF1A-EmGFP form prior to extensive egress of virus particles (Figure 8A). In Vero cells expressing KIF1A-EmGFP by plasmid transfection, without virus infection, we also observed accumulation of KIF1A-EmGFP at the tips of cell extensions (Figure 8B, arrows). Together, these latter observations indicate that these peripheral accumulations form as a result of pre-existing cellular pathways, not induced de novo by viral mechanisms.

**Figure 8.**
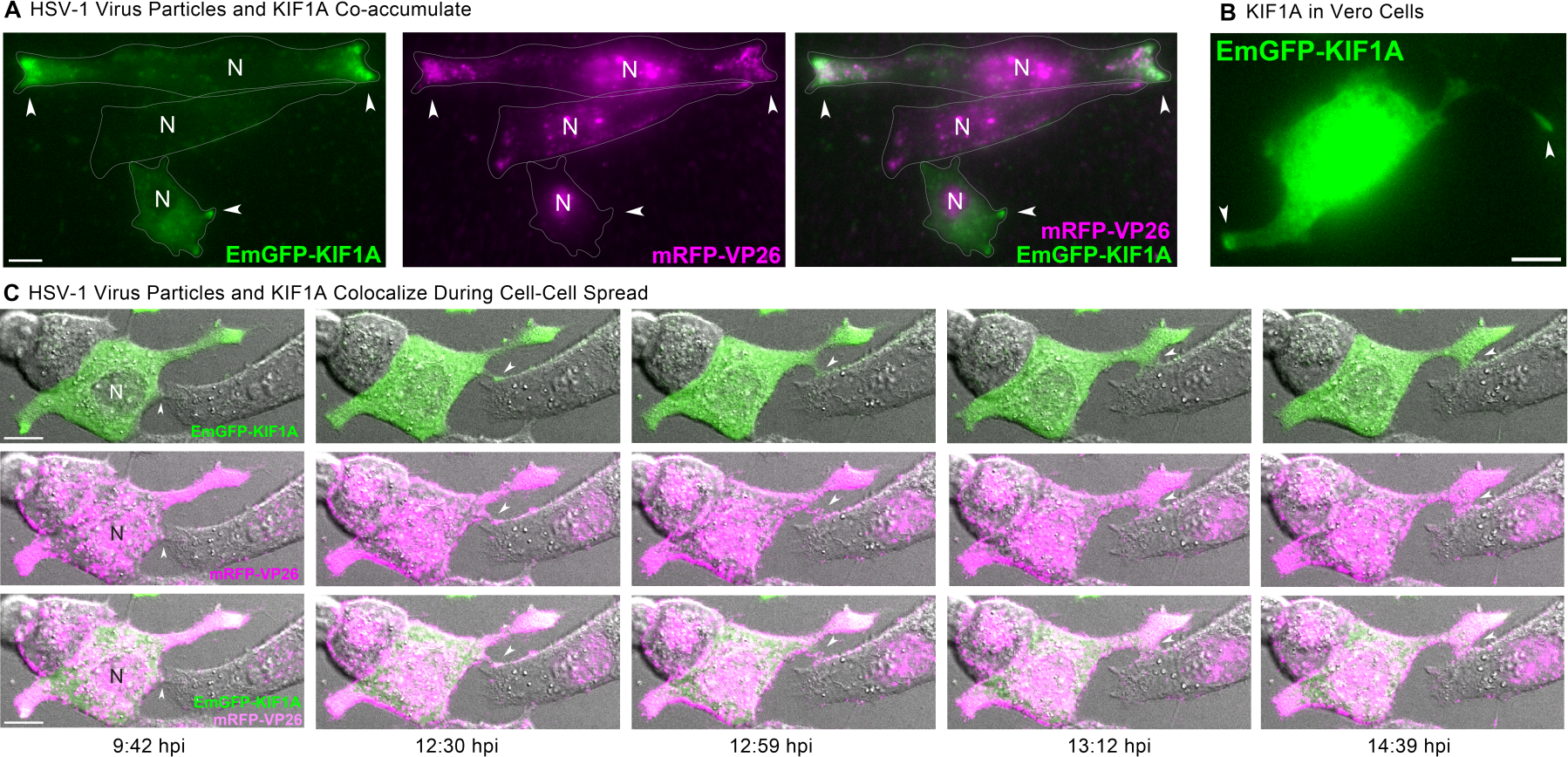
A Plus End-Directed Kinesin Motor Marks Virus Clustering Sites **A**. HSV-1 OK14 particles (magenta) co-accumulate with plus end-directed microtubule motor KIF1A (green) at the cell periphery. Vero cells were coinfected with HSV-1 OK14 and an HSV-1 amplicon vector expressing EmGFP-KIF1A, and imaged at 6 hpi. Accumulations (arrows) are indicated. **B**. In transfected cells expressing only EmGFP-KIF1A, the motor similarly accumulates at the tips of cell protrusions (arrows), in the absence of viral infection. **C**. Colocalization of EmGFFP-KIF1A and HSV-1 OK14 virus particles in cell protrusions that mediate cell-cell contact (arrows). Still images are from a timelapse movie of coinfected Vero cells imaged every 60 s from 5 hpi to 12 hpi. Scale bars represent 10*µ*m.

### Peripheral Accumulations of HSV-1 Particles Form at Cell-Cell Junctions, and Contribute to Cell-Cell Transmission

To examine the dynamics of peripheral cluster accumulation, we infected Vero cells with HSV-1, and performed timelapse confocal microscopy, imaging every 60 sec. Cells were selected for imaging at 4 hpi based on the presence of mRFP-VP26 in the cell nucleus, and then imaged overnight.In the first representative time course (Figure 9A, Movie S3), virus particles begin to form peripheral accumulations around 5 hpi (Figure 9A, arrows), including at sites of cell-cell contact. Over the next two and a half hours, these accumulations increased from a few detectable fluorescent punctae to distinct clusters of red fluorescence along these cell-cell contacts.

**Figure 9.**
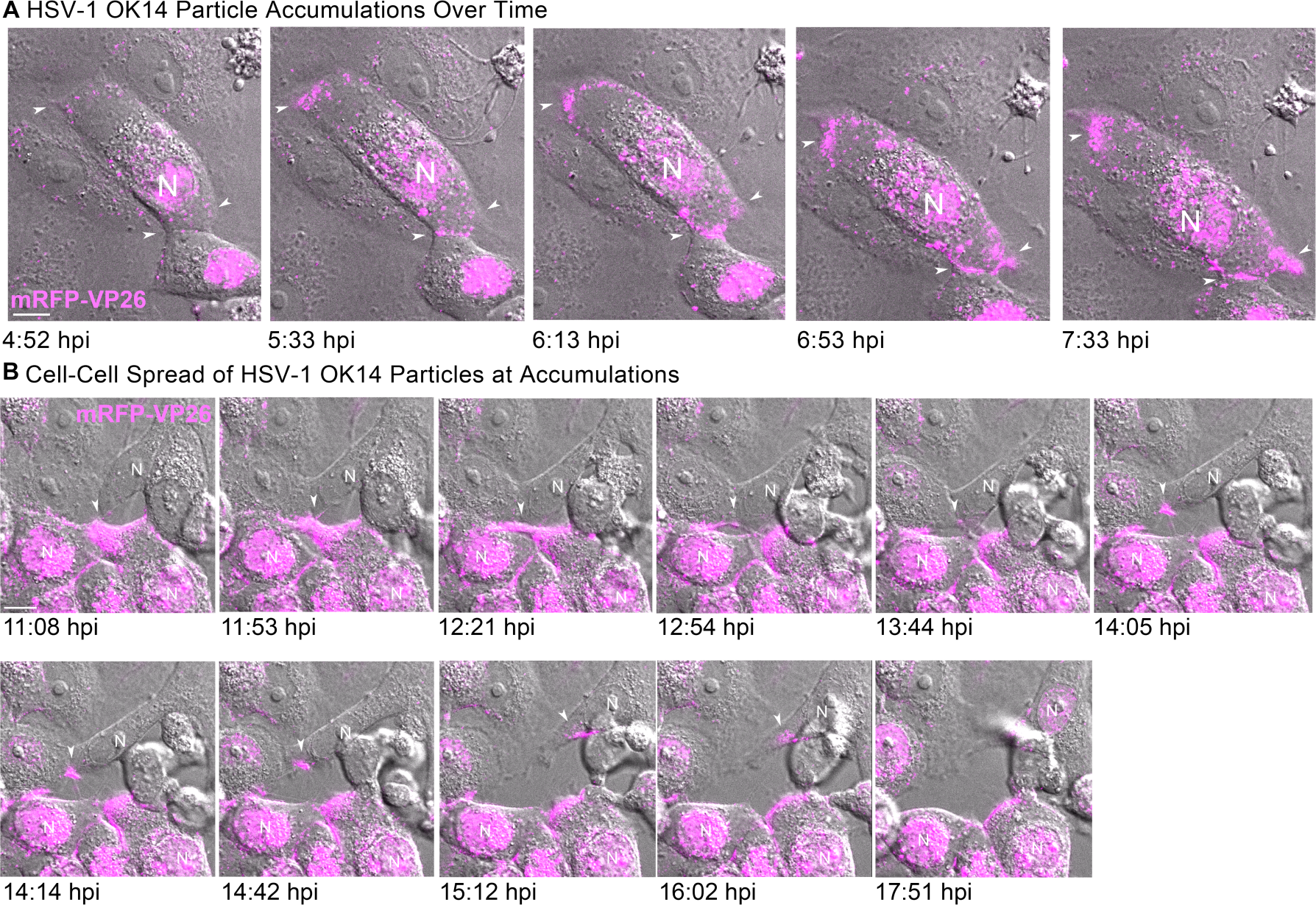
Clusters of HSV-1 Particles Form at Cell-Cell Junctions, and Contribute to Cell-Cell Transmission **A**. Representative still images from time lapse Movie S3 of virus particle accumulation in Vero cells. Cells were infected at high MOI with HSV-1 IH02 and imaged every 60 seconds from 4 hpi to 8 hpi. Arrows indicate areas of virus particle accumulation. B. Still images from Movie S4 time lapse of Vero cells infected with HSV-1 IH02 at high MOI. Images were acquired every 60 seconds from 11 hpi to 18 hpi. A large cluster of virus particles is transferred from an infected cell to an uninfected cell (arrows). This cell subsequently becomes infected, based on new capsid protein expression in the nucleus (N) at 17:51 hpi. **A-B**. Scale bars represent 10*µ*m.

In other time course examples, we observed accumulations of virus particles being transferred from one cell to another (Figure 9A-B, Movie S3-4). Because the KIF1A-EmGFP probe labels cell protrusions prior to formation of virus particle clusters, we also imaged Vero cells co-infected with HSV-1 OK14 and the KIF1A-EmGFP amplicon vector (Figure 8C). Consistent with the still images in Figure 9A-B and the timelapses of Movies S3-4, large clusters of virus particles co-accumulated with KIF1A-EmGFP in larger cell extensions. However, in timelapse imaging, we also observed the formation of smaller, more dynamic cell protrusions, which also co-accumulated virus particles and the KIF1A-EmGFP cellular marker (Figure 8C, 12:30 hpi, arrow). Over time, this smaller cell protrusion (arrows) appeared to maintain contact with a neighboring cell, with the transfer of fluorescent virus particles from cell to cell. In a second example of cell-cell transmission (Figure 9B, Movie S4), a large peripheral accumulation of virus particles extends a thin projection that transfers a bolus of many virus particles to an uninfected cell (Figure 9B, arrows). This transfer occurs at approximately 14 hpi, and the recipient cell demonstrates active viral replication about 4 hr later, based on the new appearance of mRFP-VP26 in its nucleus (Figure 9B, last image, 17:51 hpi).

These results further reinforces the idea that the large peripheral clusters of virus particles form due to preferential egress of individual virions at these locations, not by bulk release of many particles at once. In addition, these data indicate that large numbers of virions can be transferred from cell to cell via peripheral accumulations, showing that these clusters of virions can contribute to cell-cell transmission.

## 1.4 Discussion

By producing an HSV-1 recombinant virus that expresses the pH-sensitive fluorescent protein, pHluorin, we have developed a live-cell microscopy assay that allows us to visualize the process of viral egress from infected cells. pHluorin is genetically fused to the viral envelope glycoprotein gM, is incorporated into virus particles, is quenched in the lumen of cellular vesicles, but dequenches upon exocytosis, allowing detection of virus particle exocytosis. We are able to detect individual virus particles undergoing exocytosis, and while this approach had been successful in previous alpha herpesvirus studies [16, 17], this is the first time that this approach has been applied to the important human pathogen, HSV-1.

Altogether, these data show that the long-observed clustering of HSV-1 particles in Vero cells occurs due to preferential exocytosis of individual particles at these sites, rather than bulk release or post-exocytosis movements. In all the experimental conditions presented here, we have never observed bulk release of many virions at once. Our prior observations, that this clustering does not occur with PRV in PK15 cells, is the result of cell type differences, not virus differences. Both HSV-1 and PRV form clusters in Veros, primary REFs, and human cancer cell lines, but not in PK15 cells.

Differences in intracellular transport and secretory mechanisms may explain why the spatial distribution of viral egress varies across these cell types. We (and many others) have previously shown that alpha herpesvirus particles use cellular secretory pathways, regulated by Rab family GTPases, and recruit kinesin microtubule motors for intracellular transport to the site of exocytosis. Different cell types express different kinesin motors, different Rab GTPases, and other cell biological factors. In polarized epithelial cells, distinct secretory organelles sort cargoes to the apical or basolateral plasma membrane. In neurons, vesicles containing axonal cargoes can transport into the axon, but vesicles containing somatodendritic cargoes are strongly excluded from the axon. The alpha herpesviruses have evolved to modulate axonal sorting and transport by recruiting additional kinesin motors via the viral gE, gI, and US9 proteins. However, here we show that these viral factors are not required for transport to and exocytosis at preferential egress sites in these non-neuronal, non-polarized cell types.

Instead, it appears that the morphology of the microtubule cytoskeleton may be responsible for the differences in virus particle distribution during the later stages of infection. Vero cells have been documented to form distinct focal adhesion patterns [22, 74], and HSV-1 infection has been shown to alter the arrangement of microtubules [70–72, 75]. The underlying cell biological differences in cytoskeleton arrangement between different cell types, together with the effects of viral infection (i.e. microtubule stabilization) likely account for the differences we observe in the clustered/preferential egress phenotype.

## 1.5 Materials and Methods

### Cells

Vero cells (ATCC, CCL-81), PK15 cells (ATCC, CCL-33), Panc-1 (ATCC, CRL-1469) and primary rat embryonic fibroblasts (REFs) were all maintained in Dulbecco’s Modified Eagle’s Media (DMEM, Cytiva) supplemented with 10% FBS (Omega Scientific) and 1% penicillin-streptomycin (Hyclone), and incubated in a 5% CO_2_ incubator at 37°C.

REFs were collected from E16-17 Sprague-Dawley rat embryos (Charles River Laboratories), as follows: Animal work was performed in accordance with all applicable regulations and guidelines, and with the approval of the Institutional Animal Care and Use Committee (IACUC) at Arizona State University (protocol #20-1799R). The animal care and use program at Arizona State University has an assurance on file with the Office of Laboratory Animal Welfare (OLAW), is registered with the USDA, and is accredited by AAALAC International. Briefly, embryos were decapitated and internal organs removed. Remaining skin and connective tissue was trypsinized (Trypsin-EDTA, Gibco) at 37°C, pipetted vigorously in complete DMEM, and supernatants were plated onto 10cm cell culture dishes (Celltreat). REFs were passaged no more than 4 times before use in experiments [76].

### Panc-1 DHX9 knockdown cells

Lentivirus vectors expressing a pool shRNAs targeting DHX9 (shDHX9) were purchased from Santa Cruz Biotechnology. Target sequences are as follows: Sense1: CCAGAGACUUUGUUAACUAtt, Anti-sense1:UAGUUAACAAAGUCUCUGGtt, Sense2: GCAAGCGACUCUAGAAUCAtt, Antisense2: UGAUUCUAGAGUCGCUUGCtt, Sense3: GCAUGGACCUCAAGAAUGAtt, Antisense3: UCAUUCUUGAGGUCCAUGCtt. We seeded Panc-1 in a 96-well plate 24 hours prior to lentivirus transduction. Cells were treated with 5*µ*g/mL polybrene and transduced with shDHX9 lentivirus pool at MOI 10. 24 hours post-transduction, medium was exchanged with fresh media containing 10*µ*g/mL puromycin dihydrochloride, to select for stably transduced cells. Cells were maintained in media containing puromycin for 1 month before experimentation.

### Viruses

All HSV-1 or PRV strains were propagated and titered by plaque assay on Vero or PK15 cells, respectively, in DMEM supplemented with 2% FBS and 1% penicillin-streptomycin. HSV-1 17syn^+^ and OK14 were obtained from the Lynn Enquist laboratory (Princeton University) and verified by whole genome sequencing. HSV-1 OK14, which expresses an mRFP-VP26 capsid tag, was previously described [39]. HSV-1 425, which is based on HSV-1 MacIntyre and expresses an mRFP-VP26 capsid tag, was a kind gift from Esteban Engel (Princeton University) [55]. PRV 483 and PRV 495, which express gM-pHluorin and an mRFP-VP26 capsid tag, were previously described [16]. PRV BaBe was obtained from the Lynn Enquist laboratory (Princeton University) [64, 65].

### Construction of New HSV-1 Recombinants

Three confluent 10 cm dishes of Vero cells were infected with HSV 17syn^+^ at MOI of 5 pfu/cell, and incubated overnight. Infected cells were rinsed with PBS, scraped from the dish, and lysed with an NP-40/Tris buffer (140mM NaCl, 2mM MgCl2, 0.5% Nonidet P-40, 200mM Tris). Nuclei were pelleted by centrifugation, and then lysed with 1% SDS in PBS. 100*µ*g/mL proteinase K was added, and incubated at 50°C for 1 hour. DNA was then isolated by phenol-chloroform extraction and ethanol precipitation. To produce HSV-1 IH01, a shuttle plasmid containing the pHluorin coding sequence flanked by HSV-1 sequences homologous to the HSV-1 UL10/gM locus was synthesized (Genewiz). This construct was designed to insert pHluorin into the first extravirion loop of the gM protein. Vero cells were cotransfected with linearized shuttle plasmid and DNA isolated from HSV-1 17syn^+^ infected cells using JetPrime transfection reagent (Polyplus). Following reconstitution of replicating virus, plaques were screened for expression of green fluorescence and plaque purified three times. To produce HSV-1 IH02, Vero cells were co-infected with HSV-1 IH01 and OK14, and progeny were screened for red and green fluorescence, and plaque purified three times.

### Construction of New PRV Recombinants

PRV IH001 was constructed by co-infecting PK15 cells with PRV 495 and PRV BaBe. PRV 495 expresses gM-pHluorin and mRFP-VP26, but also contains a deletion in the essential UL25 gene and cannot replicate on its own. PRV BaBe contains a deletion in the US region encoding gE, gI, and US9 [64, 65]. Following co-infection, progeny plaques were screened for green and red fluorescence. Several clones were picked, plaque purified three times, and further screened for lack of gE, gI, and US9 expression via western blot (Supplemental 2).

### HSV-1 Amplicon Vector Construction and Propagation

The amplicon vector plasmid pCPD-HSV-N-EmGFP-DEST was constructed by the DNASU Plasmid Repository (Biodesign Institute, Arizona State University), as follows: The plasmid HSV-DYN-hM4Di was a gift from John Neumaier (Addgene plasmid # 53327) [77]. Unnecessary promoter and transgene sequences were removed by digestion with HindIII and religating, to produce pCPD-HSV. This plasmid contains the HSV-1 packaging signal and OriS origin of replication. The plasmid pcDNA6.2/N-EmGFP-DEST obtained from Invitrogen (ThermoFisher). The CMV promoter, Emerald GFP coding region, Gateway recombination cassette (attR1, CmR selection marker, ccdB counterselection marker, attR2), and HSV-1 TK polyadenylation signal were PCR amplified and ligated into the HindIII site on pCPD-HSV, to produce pCPD-HSV-N-EmGFP-DEST. The human KIF1A coding sequence (DNASU Plasmid Repository, #HsCD00829423, NCBI Nucleotide reference NM_001244008.2) was inserted by Gateway recombination (Invitrogen) to make an in-frame KIF1A-EmGFP fusion.

To propagate the KIF1A-EmGFP amplicon vector, 3.5 x 10^5^ HEK 293A cells were seeded into each well of a 6-well plate (Celltreat), incubated overnight, and then transfected with 3*µ*g of amplicon plasmid using Lipofectamine 2000 (Invitrogen). 24 hours after transfection, cells were infected with 10^5^ infectious units of HSV-1 OK14. Cells were incubated for another 24 hours, and then cells and supernatants were harvested. The amplicon stock was passaged at high MOI one time on Vero cells, cells and supernatants were harvested, and stored at 73×2212;80°C.

### Fluorescence microscopy

All cell types were seeded at subconfluent density (∼10^5^ cells/dish) on glass-bottom 35mm dishes (Celltreat, Ibidi, and Mattek), incubated overnight, and then infected with HSV-1 or PRV at a relatively high MOI (>1 pfu/cell). o account for differences in the efficiency of plating between different viruses and cells, the amount of inoculum needed to synchronously infect most cells was determined empirically on a case-by-case basis using fluorescence microscopy. MOIs ranged from 1-20 pfu/cell, as titered on Vero or PK15 cells for HSV-1 and PRV, respectively, without taking differences in efficiency of plating into account. HSV-1 infected cells were imaged beginning at 5-6 hpi, and PRV infected cells were imaged beginning at 4-5 hpi, unless otherwise stated. Fluorescence microscopy was performed using a Nikon Eclipse Ti2-E inverted microscope in the Biodesign Imaging Core facility at Arizona State University. This microscope is equipped with TIRF and widefield illuminators, a Photometrics Prime95B sCMOS camera, a 60X high-NA TIRF objective, and objective and stage warmers for 37°C live-cell microscopy. For widefield fluorescence, a Lumencor SpectraX LED lightsource provided 470/24nm and 550/15nm excitation for green and red fluorescent proteins, respectively.

For TIRF microscopy, 488nm and 561nm lasers were used to excite green and red fluorescent proteins, respectively. Image analysis was performed using Fiji software [78]. Fluorescence microscopy images were prepared for publication using Adjust Brightness/Contrast, Reslice (to produce kymographs), and Plot Z-axis Profile (to measure fluorescence over time) functions in Fiji. Maximum difference projections were calculated as previously described [16], using the Duplicate, Stacks->Tools, Math->Subtract, and Z Project functions in Fiji.Ensemble averages of fluorescence intensity over time in a 3×3 pixel region of interest around individual exocytosis events (Figure 3B-C, 5C, 6C) were calculated using Matlab (Mathworks).

For confocal microscopy, samples were imaged on a Nikon AX R laser scanning confocal microscope in the Biodesign Imaging Core facility (Arizona State University, Tempe, AZ) using a 60X 1.42 NA objective. EmGFP was excited at 488nm and mRFP at 568nm. Emissions for these channels were collected in the green and red ranges respectively. Images were acquired every 60 seconds with a 100ms exposure. Timelapse movies were registered using Registration->Correct 3D Drift->Correct X & Y functions in Fiji.

### Western Blot

Vero cells were infected with HSV-1 and PK15 cells were infected with PRV at high MOI and incubated overnight. Infected cells were lysed with an NP-40 lysis buffer (50mM Tris, 150mM NaCl, 1% Nonident P-40, diH2O) on ice for 3 minutes, nuclei were pelleted by centrifugation at 16000 rpm for 20 min, supernatants were mixed with 2X Laemmli sample buffer containing SDS and 2-mercaptoethanol, and heated to 95°C for 5 min. SDS-PAGE separation was run on Nu-Page precast gels (Invitrogen). Proteins were then transferred onto PVDF membrane (Immobilon-FL, Millipore) using a semi-dry transfer apparatus (BioRad). Membranes were blocked with a 5% nonfat dry milk solution, and probed with antibodies overnight. Mouse monoclonal anti-GFP antibody (Sigma) was used to detect pHluorin. Rabbit polyclonal antibodies targeting HSV-1 gM (PAS980), PRV gE, gI, and US9 were kindly provided by Lynn Enquist (Princeton University) [79, 80]. The next day, membranes were probed with fluorescent secondary antibodies (LI-COR) for one hour, washed, and imaged using a LI-COR Odyssey CLx scanner.

### Particle Imaging and pHluorin Quenching

Vero cells in 35mm cell culture dishes were infected with HSV IH02 at high MOI. 24 hours post-infection, 100*µ*L of supernatant was pipetted onto glass bottom 35mm dishes for imaging. After allowing virus particles in the supernatant to adhere non-specifically to the glass, excess media was aspirated off and HBSS (Gibco) was added to prevent drying. Virus particles were subjected to a pH change by adding 150*µ*L PBS at pH 6. pH was then returned to neutral by adding an excess of PBS at pH 8. Imaging was performed using widefield LED illumination and 60X magnification to detect individual virus particles.

### Single Step Growth Curve and Plaque Size Measurements

Vero cells were seeded to confluence in 35mm 6-well dishes and infected at MOI of 5 PFU/cell. The inoculated cells were incubated for 1 hour, washed with PBS three times, and incubated with viral medium at 37°C. At the specified time points, cells and supernatants were harvested. Mean titers were determined for each time point via serial dilution plaque assay in triplicate. Plaque sizes were measured in Fiji from brightfield and fluorescence microscopy images of 35mm wells of plaque assays from the single step growth curve.

## Statistical Analysis

Statistical tests, one-way ANOVA and Student’s T-Tests, were performed in GraphPad Prism or Microsoft Excel.

## Acknowledgements

Thank you to Joli Bastin for her work on producing the recombinant HSV IH01 and IH02 strains carrying gM-pHluorin.

